# Gains through selection for grain yield in a winter wheat breeding program

**DOI:** 10.1101/734194

**Authors:** Dennis N. Lozada, Arron H. Carter

## Abstract

Increased genetic gains for complex traits in plant breeding programs can be achieved through different selection strategies. The objective of this study was to compare potential gains for grain yield in a winter wheat breeding program through estimating response to selection *R* values across several selection approaches including phenotypic (PS), marker-based (MS), genomic (GS), and a combination of PS and GS. Five populations of Washington State University (WSU) winter wheat breeding lines evaluated from 2015 to 2018 in Lind and Pullman, WA, USA were used in the study. Selection was conducted by selecting the top 20% of lines based on observed yield (PS strategy), genomic estimated breeding values (GS), presence of yield “enhancing” alleles of the most significant single nucleotide polymorphism (SNP) markers identified from genome-wide association mapping (MS), and high observed yield and estimated breeding values (PS+GS). Overall, PS compared to other individual strategies showed the highest response. However, when combined with GS, a 23% improvement in *R* for yield was observed, indicating that gains could be improved by complementing traditional PS with GS. Using GS alone as a selection strategy for grain yield should be taken with caution. MS was not that successful in terms of *R* relative to the other selection approaches. Altogether, we demonstrated that gains through increased response to selection for yield could be achieved in the WSU winter wheat breeding program by implementing different selection strategies either exclusively or in combination.

## Introduction

The challenge to develop higher yielding, climate resilient, disease- and pest-resistant, and more nutritious crops has never been more urgent considering the anticipated population growth in the next 30 years [1]. As such, improving genetic gains or performance for important traits such as yield, disease resistance, and adaptation in staple crops such as wheat (*Triticum aestivum* L.) has been the goal of many breeding programs. Genetic gain is the predicted change in mean value of a trait within a population under selection [2] and is represented by what is more commonly known as the “breeder’s equation” [3]. To increase genetic gains, an increase in the phenotypic variability, accuracy of selection, and selection intensity, or a decrease in generation time for cultivar development, is necessary [4]. Phenotypic, genomic, and marker-based selection approaches could be used to increase either of the factors mentioned to achieve improved gains.

In bread wheat, phenotypic selection for superior genotypes, characterized primarily by a “non-shattering” phenotype, begun during its domestication [5]. This “unconscious” breeding resulted from the unintentional selection of lines that were more adapted and productive under early farming practices and by natural selection in the fields [6]. The “empirical” and “scientific” breeding followed the “unconscious”, which resulted in the development of wheat lines with improved characteristics in breeding programs [6]. Currently, plant breeders have access to advanced genome and phenotypic-based selection strategies to fast-track genetic improvement and increase gains for key traits in wheat [1].

Several studies evaluated the gains which could be achieved by applying different selection strategies particularly for increasing resistance to specific diseases in wheat. Rutkoski et al. [7] compared gains for phenotypic and genomic selection for quantitative stem rust resistance and observed that genomic selection could perform as well as phenotypic selection for stem rust resistance improvement but can result in less genetic variation over time. Significant gains using marker-assisted selection for Fusarium head blight (FHB) resistance were also observed in the University of Minnesota wheat breeding program due to the presence of a major quantitative trait locus. Using closely linked and diagnostic markers for *Fhb1* caused a 27% reduction in disease symptoms throughout the breeding programs [8]. In another study, FHB severity in winter wheat was reduced by 6 and 5% using phenotypic and marker-aided selection, respectively [9]; whereas marker-assisted breeding for severity and deoxynivalenol (DON) content resulted in higher gains on an annual basis in spring wheat [10]. Both studies observed a large variation for FHB resistance in the marker-selected lines demonstrating the need to complement marker selection with phenotypic selection to further enhance gains.

Grain yield is a complex trait controlled mainly by many loci with small effects [11–13] and this makes yield more difficult to examine than disease resistance. Improvement in grain yield, however, remains the prime emphasis of many wheat breeding programs [14], and with that, it is necessary to measure gains achieved through different breeding and selection strategies. Given that there are several selection approaches used in plant breeding, we were interested in quantifying the possible gains for grain yield which could be attained when these methods are implemented alone or in combination with others in a winter wheat breeding program. The objective of this study was to compare the projected gains for yield resulting from using different selection strategies in the Washington State University (WSU) winter wheat breeding program. Empirical datasets for grain yield collected from over 2,200 WSU winter wheat breeding lines grown from 2015 to 2018 were evaluated. The different selection strategies assessed included phenotypic, marker, genomic, and a combination of phenotypic and genomic selection. Potential gains for yield represented as the response to selection *R* were calculated for these selection strategies.

## Materials and methods

### Winter wheat populations

A total of five different populations of soft winter wheat lines adapted to the US Pacific Northwest was used in the study. These populations included an association mapping panel (AMP), two F5, and two double haploid (DH) populations of WSU winter wheat breeding lines. The AMP consisted of 456 lines evaluated in Lind (LND) and Pullman (PUL) WA, USA between 2015 and 2018. Significant soil crusting delayed the growth of the winter wheat lines in LND in 2016 and hence the AMP was not evaluated for this site-year. The F5 lines comprised of 61 and 501 lines planted in 2017 in LND (LND17_F5) and PUL (PUL17_F5), WA respectively. The DH panels were evaluated in LND and PUL in 2018 and consisted of 447 (LND18_DH) and 759 (PUL18_DH) winter wheat breeding lines.

### Phenotypic data collection and analyses

Grain yield (in t ha ^-1^) was assessed by harvesting whole plots using a Zurn^®^ 150 combine (Waldenburg, Germany). Adjusted yields were calculated under an augmented design with replicated checks and un-replicated genotypes on each block through the Augmented Complete Block Design (ACBD) in R program [15]. The winter wheat line ‘Eltan’ [16] was used as a check in LND, and ‘Madsen’ [17] was used as a check in PUL for the 2015-2018 growing seasons for the AMP. Checks for the LND17_F5 included the lines ‘Bruehl’ [18], Eltan, ‘Otto’ [19], ‘Jasper’ [20], Madsen, and ‘Xerpha’[21], whereas ‘Brundage’[22], Jasper, Madsen, ‘Puma’[23], ‘UI Bruneau’, and ‘Xerpha’ were used for the PUL17_F5 population. Jasper, Otto, and Xerpha were used as checks for LND18_DH; whereas Jasper, Madsen, Puma, and Xerpha were used as checks for the PUL18_DH panel.

Adjusted values for yield were calculated employing two statistical models following Lozada and Carter [24]. Briefly, the models used were:

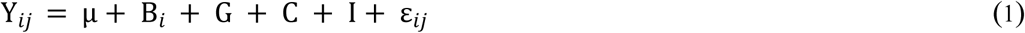

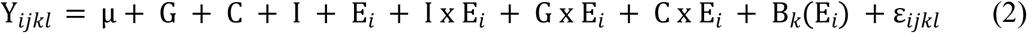

where Y is the trait of interest; µ is the effect of the mean; B_i_ is the effect of the *i*th block; G corresponds to the un-replicated genotypes; C is the effect of the replicated checks on each block; E_i_ is the effect of the *i*th environment; I is the effect of the identifier of the checks; I x E_i_, G x Ei, and C x E_i_ are the effects of check identifier by environment, genotype by environment, and check by environment interactions, respectively; B_k_(E_i_) is the effect of block nested within each environment; and ε is the standard normal error [15]. Best linear unbiased estimates (BLUEs) were calculated for individual environments (eq. (1)), whereas best linear unbiased predictors (BLUPs) were computed for the combined analyses across locations (eq. (2)). Factors were considered fixed when calculating BLUEs whereas effects were regarded as random for calculating BLUPs.

### Genome-wide association study and genomic predictions

SNP genotyping was conducted using genotyping-by-sequencing (GBS) [25] through the NC State University Genomics Sciences Laboratory in Raleigh, NC, USA. The restriction enzymes *MspI* and *PstI* were used for GBS. SNPs were filtered for minor allele frequency (MAF) of > 0.05 and 10% missing data. After quality control, 16,233 markers (genotype data 1, GD1; S1 File) remained and were used for genome-wide association study (GWAS) using a fixed and random effects circulating probability unification (FarmCPU) [26] kinship model in R [27]. SNP loci were declared to be significant under a Benjamini-Hochberg false discovery rate (FDR) [28] threshold of 0.05. The percent phenotypic variation explained (R^2^) by each significant SNP locus was calculated using a stepwise regression model in JMP^®^ Genomics v.8.1 [29], where the R^2^ value when a marker was removed from the regression model was subtracted from the total R^2^ when all the significant SNPs were fitted in the model.

Genomic predictions and genomic estimated breeding value (GEBV) calculations were implemented in the iPAT (Intelligent Prediction and Association Tool) package [30], where a ridge regression best linear unbiased prediction (RRBLUP) selection model [31] was trained using the AMP to predict the yield performance of WSU F5 and DH winter wheat breeding lines for independent validations. This prediction model considers markers to have effects toward zero with a common variance [31]. RRBLUP uses the ‘mixed.solve’ function in the form: **y** = **Xβ** + **Zu** + ε, **u** ∼ N (0, **K**σ^2^ _*u*_), where **X** is a full-rank design matrix for the fixed effects, **β**; **Z** is the design matrix for the random effects **u**, **K** is a semidefinite covariance matrix, obtained from markers using the ‘A.mat’ (additive relationship matrix function); residuals are normal with a mean of zero and constant variance; and **u** and ε independent [31].

A total of 11,089 high-quality GBS-derived SNP markers common to both the AMP and the validation sets (genotype data 2, GD2; S2 File) were used for genomic predictions. GD2 was a subset of GD1 which was used to perform association analyses using the AMP. Phenotypic data for yield in the validation populations (i.e. the F5 and DH breeding lines) were masked by representing them as “NA” during each analysis. Predictive ability for the independent validations were calculated as the Pearson correlation between GEBV and adjusted yield for the F5 and DH wheat breeding lines. For the GWAS-assisted GS, the top five most significant SNPs based on an FDR of 0.05 were fitted in an RRBLUP genomic prediction model as fixed effects in iPAT. A total of seven BLUE and two BLUP yield datasets were used for GWAS and genomic predictions. Relatedness between the diversity training panel and winter wheat test lines were assessed using Rogers genetic distances calculated in JMP Genomics v.8.0.

### Correlation between GEBV for one year and observed yield in the succeeding year

The relationships between calculated breeding values for one year and its corresponding adjusted yield on the succeeding year were evaluated by calculating GEBV of the lines in the AMP and comparing them to their adjusted yield in the next growing season (e.g. GEBV for PUL2015 was compared to adjusted yield in PUL2016). GEBVs were calculated by performing a five-fold cross-validation for the AMP, where 80% of the lines were used to predict the remaining 20% using an RRBLUP model in iPAT for the GS1 scenario. The Pearson correlation coefficients between GEBV and adjusted yield were calculated.

### Selection strategies and response to selection

Different selection approaches for grain yield, namely phenotypic (PS), marker-based (MS), genomic (GS), and phenotypic + genomic (PS+GS) selection were compared in this study. For PS, the top 20% of the F5 and DH lines based on adjusted values for yield were selected. In MS, lines having five yield “enhancing” loci identified from association mapping using the AMP were selected. These loci represented the five most significant SNPs based on a Benjamini-Hochberg FDR of 0.05 across datasets. In the GS approach, the top 20% of the breeding lines having the highest GEBV were identified through independent predictions by training the AMP to predict yield of the F5 and DH breeding lines (GS1). In another GS scenario, five of the most significant markers identified from association mapping using the AMP were included in the selection model as fixed effects to predict yield for the breeding lines using an RRBLUP model (GS2). Finally, for the PS+GS approach, lines having the top 20% highest adjusted grain yield and the highest GEBV were selected for both GS1 (PS+GS1) and GS2 (PS+GS2). The average of the adjusted yield of the corresponding lines selected for each of the selection strategy was reported. Comparisons between mean yield achieved by applying the different selection approaches were also compared to the mean of the check lines.

Gains achieved through each selection approach were represented as the response to selection, *R*, calculated as *R*= *H*^*2*^S [32], where *H*^*2*^ is the broad-sense heritability calculated as 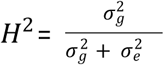 and *S* is the selection differential, calculated as *S*= µ_Selected_-µ_Unselected_, where µ_Selected_ is the mean yield for the lines with a selection strategy implemented and µ_Unselected_ is the mean yield of the lines without selection applied [33].

## Results

### Significant marker-trait associations

A total of 24 significant marker-trait associations (MTAs) distributed across 14 chromosomes were identified for yield in the AMP under a kinship model and an FDR of 0.05 (Table 1). Three of these grain yield-related MTAs were on chromosomes 3B and 5B. The percent variation explained by each significant marker ranged between 0.001 (*S2B_239862383*) and 0.05 (*S7A_545581556*) identified in LND17 and PUL15, respectively. No SNP locus was identified to be significant across multiple datasets. FDR adjusted *P*-values for the significant markers ranged between 6.43E-06 (*S1A_535858090*) and 0.048 (*S3B_482345832*), whereas allele effects ranged between -0.39 and 0.26. The significant MTAs had an average minor allele frequency of 0.32.

**Table 1.** SNP markers associated with grain yield identified in a diverse training panel of US Pacific Northwest winter wheat lines (*N*= 456 lines).

| SNP | Dataset | Chr. | Position (bp) | FDR adj. *P*-value <sup>a</sup> | Minor allele frequency | Phenotypic variation explained, R<sup>2</sup> |
| --- | --- | --- | --- | --- | --- | --- |
| **S1A_497083519** | PUL15 | 1A | 497083519 | 0.01 | 0.38 | 0.0213 |
| **S1A_535858090 <sup>b</sup>** | **PUL18** | **1A** | **535858090** | **6.43E-06** | **0.34** | **0.0324** |
| **S1B_8150831** | PUL18 | 1B | 8150831 | 0.01 | 0.14 | 0.0456 |
| **S2A_752287563** | LND17 | 2A | 752287563 | 0.02 | 0.36 | 0.0129 |
| **S2B_239862383** | LND17 | 2B | 239862383 | 0.01 | 0.37 | 0.0001 |
| **S2B_775486161** | PUL18 | 2B | 775486161 | 0.01 | 0.41 | 0.0175 |
| **S2D_639821303** | LND17 | 2D | 639821303 | 0.03 | 0.18 | 0.0154 |
| **S2D_642029978** | LND17 | 2D | 642029978 | 0.02 | 0.08 | 0.0007 |
| **S3A_22831895** | LND18 | 3A | 22831895 | 0.04 | 0.42 | 0.0241 |
| **S3A_567971108** | PUL15 | 3A | 567971108 | 0.009 | 0.19 | 0.0145 |
| **S3B_482345832** | PUL18 | 3B | 482345832 | 0.05 | 0.47 | 0.0025 |
| **S3B_561570016** | **PUL18** | **3B** | **561570016** | **0.003** | **0.26** | **0.0159** |
| **S3B_818284683** | **PUL15** | **3B** | **818284683** | **0.005** | **0.47** | **0.0217** |
| **S3D_325690** | LND17 | 3D | 325690 | 0.01 | 0.21 | 0.0033 |
| **S5B_29125444** | LND17 | 5B | 29125444 | 0.006 | 0.08 | 0.0005 |
| **S5B_47592949** | PUL18 | 5B | 47592949 | 0.01 | 0.31 | 0.0090 |
| **S5B_679577399** | LND18 | 5B | 679577399 | 0.04 | 0.32 | 0.0189 |
| **S6A_601959488** | LND17 | 6A | 601959488 | 0.04 | 0.21 | 0.0012 |
| **S6B_118986455** | **LND18** | **6B** | **118986455** | **0.0001** | **0.15** | **0.0286** |
| **S6B_33331876** | PUL18 | 6B | 33331876 | 0.014 | 0.50 | 0.0008 |
| **S7A_545581556** | PUL15 | 7A | 545581556 | 0.03 | 0.46 | 0.0507 |
| **S7A_61774265** | **PUL15** | **7A** | **61774265** | **0.001** | **0.36** | **0.0228** |
| **S7B_711208053** | PUL15 | 7B | 711208053 | 0.007 | 0.45 | 0.0205 |
| **S7D_635365239** | PUL15 | 7D | 635365239 | 0.02 | 0.46 | 0.0103 |

<sup>a</sup> FDR- False discovery rate
<sup>b</sup> Significant SNPs highlighted in bold text were included in the prediction model as fixed effects for a GWAS-assisted genomic selection scenario

### Predictive ability and genomic estimated breeding values for grain yield

Prediction ability for the GS1 scenario under independent validations were low, ranging from -0.21 (PUL16 predicting LND17_F5) to 0.21 (PUL15 predicting LND17_F5) across the wheat breeding lines (Fig 1). Overall, higher accuracies were observed for predicting the F5 lines compared with the DH populations (0.03 vs. 0.0002). No significant differences were observed for accuracies when models were trained using the LND and PUL datasets (0.01 vs. 0.02). Predicting LND17_F5 and LND18_DH wheat breeding lines using LND datasets resulted in a mean prediction ability of -0.01 whereas using PUL17_F5 and PUL18_DH as validation populations resulted in a mean predictive ability of 0.01. Across environment predictions using the LND yield datasets to predict PUL17_F5 and PUL18_DH populations resulted in a mean of 0.04, whereas using PUL datasets to predict LND17_F5 and LND_18 DH resulted in a mean of 0.02. BLUP datasets showed an advantage over BLUE datasets for predictions (0.02 vs. 0.01) across different validation populations. Mean grain yield GEBV for all the breeding lines across each dataset ranged between 2.22 (LND15 as training dataset) and 9.99 (PUL18 as training dataset) for GS1 (S1 Table).

**Fig 1.**
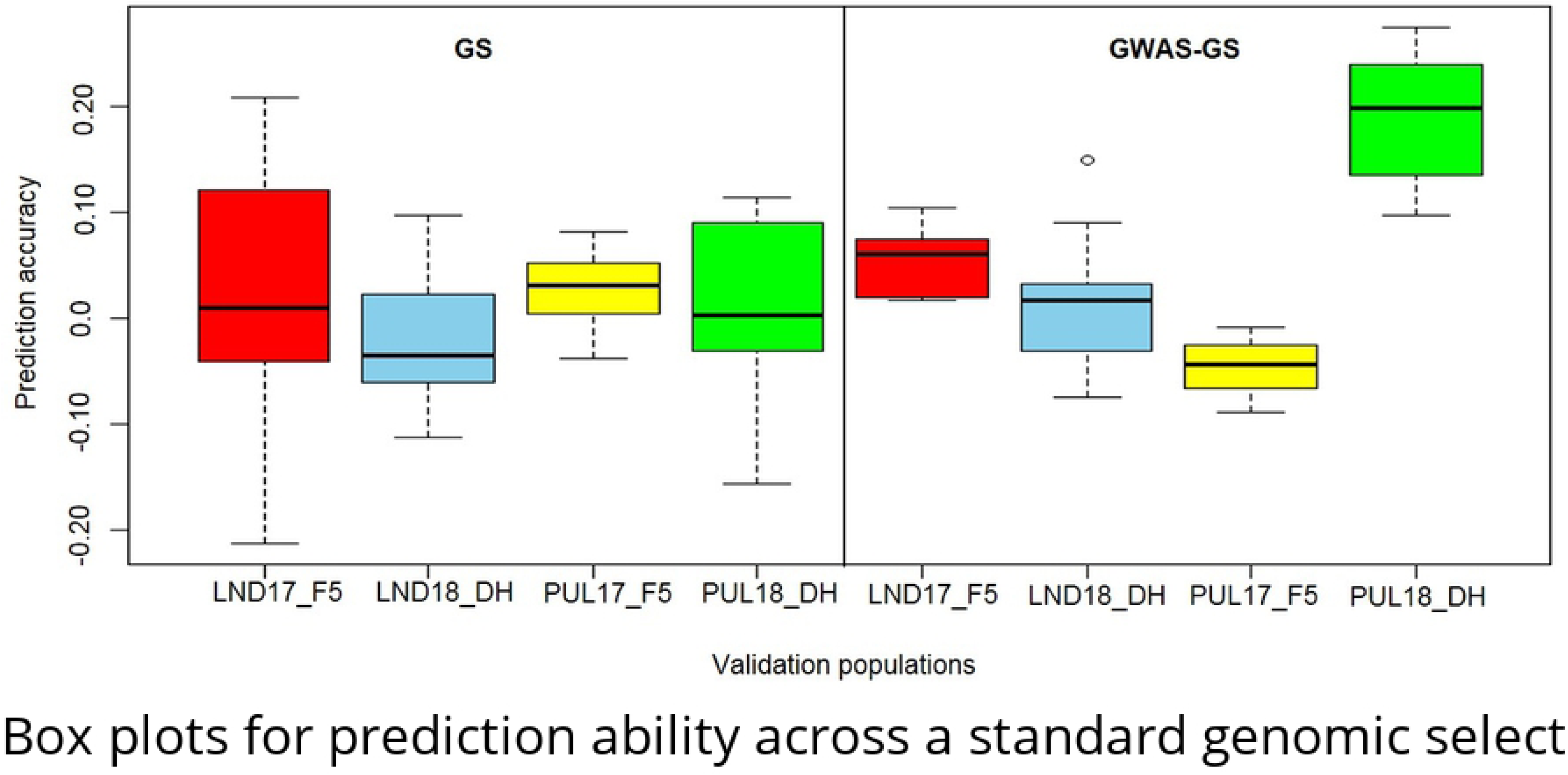
Box plots for prediction ability across a standard genomic selection approach using RRBLUP (GS1) and a GWAS-assisted GS scheme (GS2) for grain yield in a winter wheat breeding program.

Predicting grain yield using SNPs identified from GWAS as fixed effects in the model (GS2) did not result in significant differences in mean accuracy overall, although it resulted in an increase in predictive ability (0.05 vs. 0.02). Significant differences (*P <* 0.05) for mean prediction ability, nonetheless, were observed for PUL17_F5 and PUL18_DH. Prediction ability for GS2 ranged between -0.09 (PUL18 predicting PUL17_F5) and 0.27 (PUL15 predicting PUL18_DH). Highest mean prediction ability across datasets was observed for PUL18_DH (0.19), followed by LND17_F5 (0.05), LND18_DH (0.02), and PUL17_F5 (−0.04). Predicting yield using BLUP datasets did not give advantage over to using BLUEs for predictions. In contrast to GS1, within environment predictions resulted in a 50% gain in mean prediction ability compared to predicting across environments. Similar with the GS1 scenario, the highest mean GEBV for yield was observed for PUL18 (7.68) whereas the lowest was observed for LND15 (1.74) (S2 Table).

Correlations between GEBV and adjusted yield for the winter wheat breeding lines were low to high, ranging between 0.08 (LND18) and 0.71 (PUL_Com). Scatterplots showing positive significant (*P* < 0.0001) relationships between breeding values and adjusted yield for LND15, LND_Com, PUL16 and PUL17 are shown in Fig 2. Likewise, significant associations (*P* < 0.0001) between GEBV and yield were observed across growing seasons for the diverse population of US PNW winter wheat lines (AMP) (Fig 3). Correlation coefficients ranged from 0.003 (PUL15GEBV_PUL16GY) to 0.22 (PUL17GEBV_PUL18GY).

**Fig 2.**
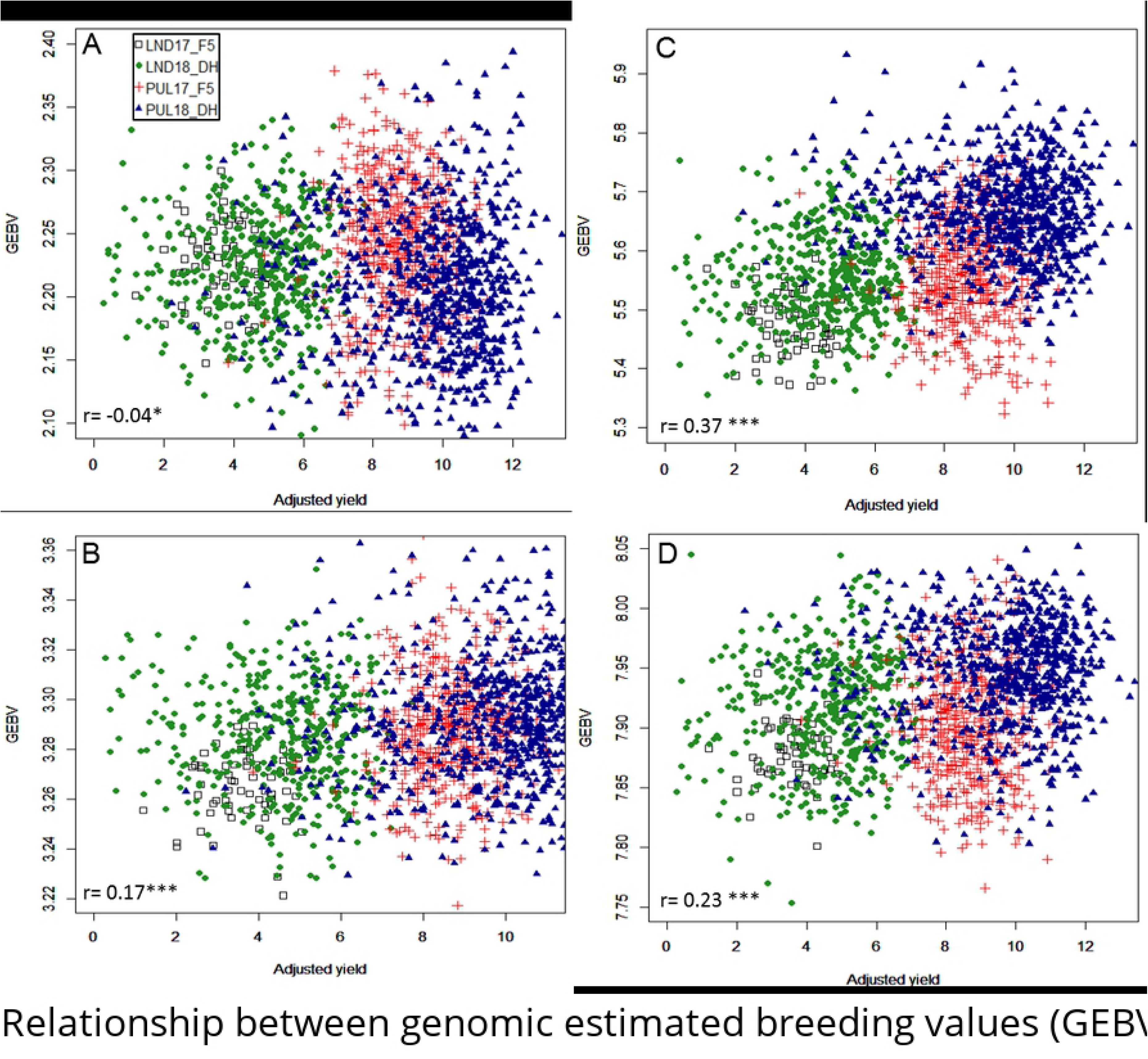
Relationship between genomic estimated breeding values (GEBV) and adjusted yield (in t ha ^-1^) for the F5 and DH wheat breeding lines for (A) LND15; (B) LND_Com; (C) PUL16; and (D) PUL17 training population datasets. **\***-Significant correlation coefficient at *P* < 0.05; ***-significant correlation at *P* < 0.0001.

**Fig 3.**
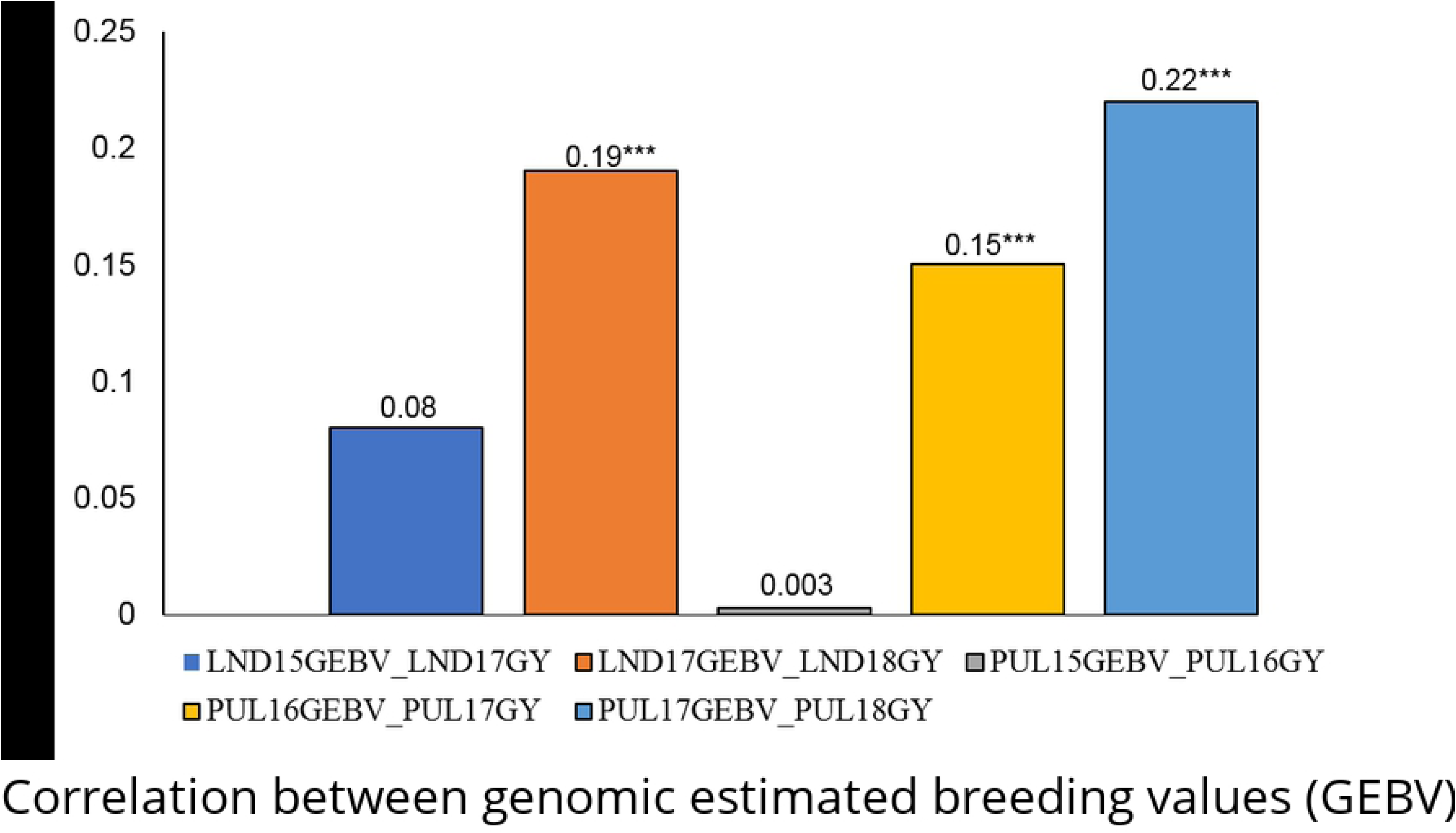
Correlation between genomic estimated breeding values (GEBV) and adjusted yield for consecutive growing seasons for a diverse association mapping population (AMP) of US Pacific Northwest winter wheat evaluated in Lind (LND) and Pullman (PUL), WA from 2015-2018. ***-Significant correlation at *P* < 0.0001

### Response to selection across different selection strategies

The highest average value for response to selection, *R*, was highest for PS, being the baseline method (1.61) (Table 2). Negative mean values for selection response were observed for both GS1 (−0.003) and MS (−0.35) (Tables 2 and 3). No line was selected under the LND17_F5 population using an MS approach, whereas there were four, 86, and 11 lines containing five favorable alleles for the most significant SNPs identified from GWAS for LND18_DH, PUL17_F5, and PUL18_DH, respectively. Using both PS+GS1 and PS+GS2 strategies, with mean *R* of 0.63 and 0.53 respectively, were more advantageous in terms of response than MS, GS1, and GS2, (Table 4). Using GWAS-derived SNP markers as fixed effects in the prediction model in the GS2 scenario resulted in higher mean *R* (0.10) compared to GS1 (−0.003). The number of lines selected on both PS and GS ranged from 0 to 44 for both PS+GS1 and PS+GS2 approaches. There were no breeding lines selected for both PS and GS scenarios when PUL16 was used to predict LND17_F5. There were 16 values for *R* (44%) for the PS+GS1 that were greater than the *R* value using the PS alone. On the other hand, only 13 *R* values (36%) for the PS+GS2 were greater than the *R* for PS (Table 4, underscored and boldfaced values).

**Table 2.** Response to selection, *R* based on phenotypic selection (PS) and marker-based selection (MS) for grain yield in US Pacific Northwest winter wheat.

| Test population | No. of lines selected <sup>a</sup> | Pop. mean (without selection) | Mean (with selection) | Selection differential <sup>b</sup> | $H^2$ <sup>c</sup> | Response to Selection <sup>d</sup> |
| --- | --- | --- | --- | --- | --- | --- |
| **<u>PS</u>** |  |  |  |  |  |  |
| **LND17_F5** | 12 | 3.58 | 4.67 | 1.09 | 0.15 | 0.16 |
| **LND18_DH** | 90 | 4.57 | 6.32 | 1.75 | 0.56 | 0.98 |
| **PUL17_F5** | 100 | 8.66 | 10.10 | 1.44 | 0.13 | 0.19 |
| **PUL18_DH** | 150 | 9.62 | 11.68 | 2.06 | 0.53 | 1.09 |
| **<u>MS</u>** |  |  |  |  |  |  |
| **LND17_F5** | 0 | 3.58 | NA | NA | 0.15 | NA |
| **LND18_DH** | 4 | 4.57 | 4.19 | -0.38 | 0.56 | -0.21 |
| **PUL17_F5** | 86 | 8.66 | 8.52 | -0.14 | 0.13 | -0.02 |
| **PUL18_DH** | 11 | 9.62 | 8.09 | -1.53 | 0.53 | -0.81 |

<sup>a</sup> Number of lines selected based on selecting the top 20% of lines on each test population based on adjusted yield values (for PS); and based on the mean yield of lines having yield “enhancing” SNPs identified through an association mapping approach using an independent population of winter wheat lines (for MS)
<sup>b</sup> Calculated as the difference between the mean yield of lines with selection and mean yield without selection, $S = \mu_{\text{Sel}} - \mu_{\text{Unselected}}$
<sup>c</sup> Broad-sense heritability
<sup>d</sup> Calculated as $R = H^2S$

**Table 3.**
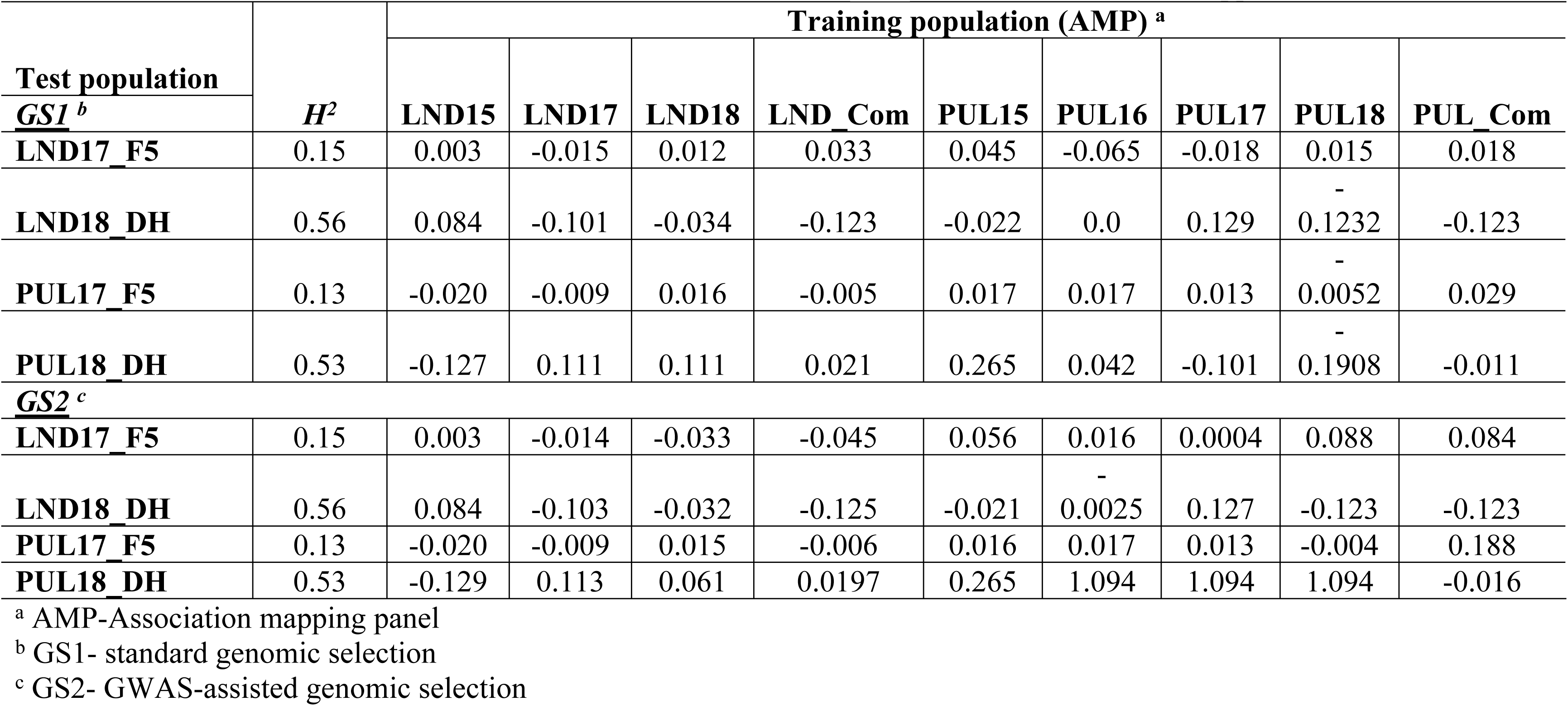
Response to selection, *R* for GEBV-based selection (GS1 and GS2) strategies for grain yield in US Pacific Northwest winter wheat. *R* values calculated based on the mean of population without selection applied.

**Table 4.**
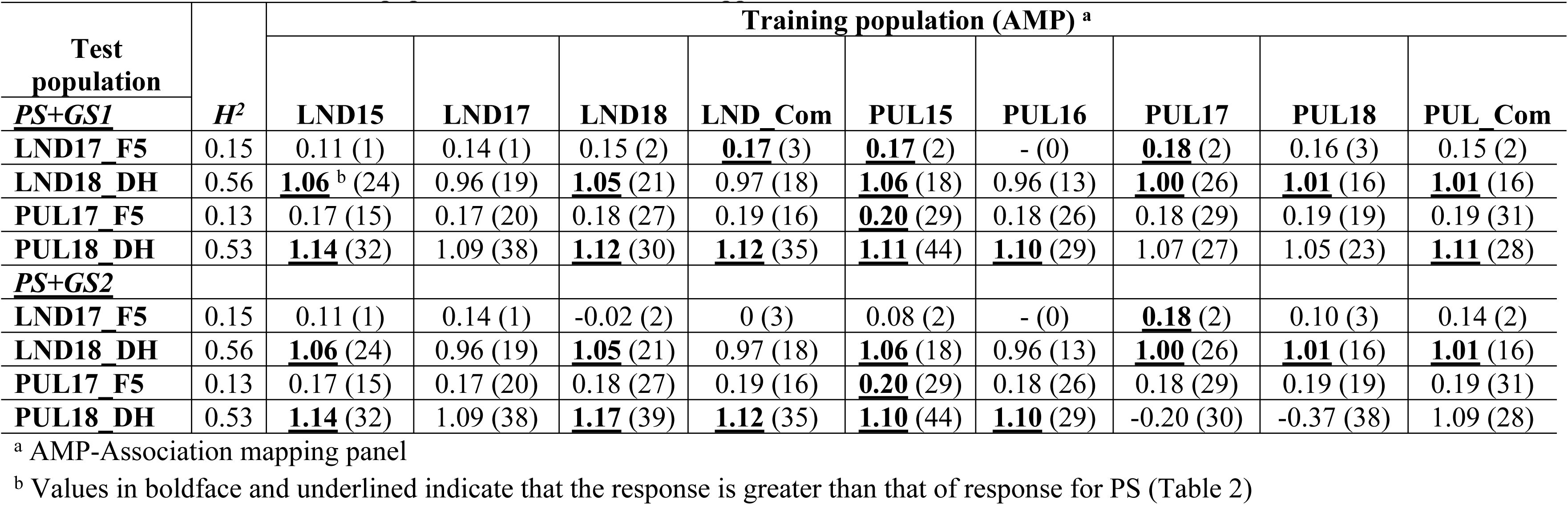
Response to selection, *R*, for phenotypic + genomic (PS+GS1 and PS+GS2) selection strategies and number of lines selected in combining both approaches for selection (in parentheses) of yield in US Pacific Northwest winter wheat. *R* values calculated based on the mean of population without selection applied.

| Test population | $H^2$ | Training population (AMP) <sup>a</sup> | | | | | | | | |
| --- | --- | --- | --- | --- | --- | --- | --- | --- | --- | --- |
|  |  | LND15 | LND17 | LND18 | LND_Com | PUL15 | PUL16 | PUL17 | PUL18 | PUL_Com |
| **<u>PS+GS1</u>** |  |  |  |  |  |  |  |  |  |  |
| LND17_F5 | 0.15 | 0.11 (1) | 0.14 (1) | 0.15 (2) | **<u>0.17</u>** (3) | **<u>0.17</u>** (2) | - (0) | **<u>0.18</u>** (2) | 0.16 (3) | 0.15 (2) |
| LND18_DH | 0.56 | **<u>1.06</u>** <sup>b</sup> (24) | 0.96 (19) | **<u>1.05</u>** (21) | 0.97 (18) | **<u>1.06</u>** (18) | 0.96 (13) | **<u>1.00</u>** (26) | **<u>1.01</u>** (16) | **<u>1.01</u>** (16) |
| PUL17_F5 | 0.13 | 0.17 (15) | 0.17 (20) | 0.18 (27) | 0.19 (16) | **<u>0.20</u>** (29) | 0.18 (26) | 0.18 (29) | 0.19 (19) | 0.19 (31) |
| PUL18_DH | 0.53 | **<u>1.14</u>** (32) | 1.09 (38) | **<u>1.12</u>** (30) | **<u>1.12</u>** (35) | **<u>1.11</u>** (44) | **<u>1.10</u>** (29) | 1.07 (27) | 1.05 (23) | **<u>1.11</u>** (28) |
| **<u>PS+GS2</u>** |  |  |  |  |  |  |  |  |  |  |
| LND17_F5 | 0.15 | 0.11 (1) | 0.14 (1) | -0.02 (2) | 0 (3) | 0.08 (2) | - (0) | **<u>0.18</u>** (2) | 0.10 (3) | 0.14 (2) |
| LND18_DH | 0.56 | **<u>1.06</u>** (24) | 0.96 (19) | **<u>1.05</u>** (21) | 0.97 (18) | **<u>1.06</u>** (18) | 0.96 (13) | **<u>1.00</u>** (26) | **<u>1.01</u>** (16) | **<u>1.01</u>** (16) |
| PUL17_F5 | 0.13 | 0.17 (15) | 0.17 (20) | 0.18 (27) | 0.19 (16) | **<u>0.20</u>** (29) | 0.18 (26) | 0.18 (29) | 0.19 (19) | 0.19 (31) |
| PUL18_DH | 0.53 | **<u>1.14</u>** (32) | 1.09 (38) | **<u>1.17</u>** (39) | **<u>1.12</u>** (35) | **<u>1.10</u>** (44) | **<u>1.10</u>** (29) | -0.20 (30) | -0.37 (38) | 1.09 (28) |

<sup>a</sup> AMP-Association mapping panel
<sup>b</sup> Values in boldface and underlined indicate that the response is greater than that of response for PS (Table 2)

Significant differences were observed between the mean *R* values for PS, GS, and MS when the mean of the checks was compared to the mean yield for the population under selection (S1-S5 Tables). Mean *R* values for PS and PS+GS1 both resulted in a 56% gain in response when compared to the mean of the checks. A total of 16 selection response values (44%) for the PS+GS1 showed higher *R* compared to the PS, whereas no *R* value for the PS+GS2 was observed to be greater than that for PS alone (S5 Table). Likewise, for the other selection strategies, an improvement in *R* was observed when check means were used to calculate selection responses (S3-S5 Tables).

## Discussion

This study reports the potential gains, represented as the response to selection *R*, which could be achieved through employing different selection strategies for grain yield in a winter wheat breeding program. Among the selection strategies evaluated were phenotypic (PS), marker-based (MS), genomic (GS), and the combination of PS and GS (PS+GS) under independent predictions.

Phenotypic selection (PS) showed an advantage over the other selection approaches in terms of *R*. Selecting a portion of lines (i.e. top 20%) based only on the adjusted yield for the F5 and DH wheat breeding lines showed a 24% gain on yield relative to the mean of the unselected population. Being the reference selection approach used in the current study, *R* values for all the other strategies should only be less than or equal to the *R* for PS. Nevertheless, it was observed that combining PS with different GS approaches (PS+GS1 and PS+GS2) under independent predictions for some of the datasets resulted in improved *R* relative to that of the PS. This indicates the potential of achieving increased gains when selecting for lines having high observed yield and high estimated breeding values (GEBV). Therefore, when performing selections, breeders could consider both information from PS and GS (through GEBV) to select lines with improved yield potential which could result in increased gains. Selecting entries having high observed yield and high breeding values could give an opportunity to choose lines that are likely to do well across environments and years in comparison to lines selected based on phenotype alone in a single year [34]. One caveat for using the PS+GS approach for selection, however, is that in some instances, there would be no lines that have both high GEBV and high observed yield selected, as in the case of using PUL16 dataset for predictions. This issue could be circumvented by evaluating more lines and increasing the selection intensity in the breeding program which could improve the chances of selecting lines having high phenotypic value and high GEBV.

Predictive ability for grain yield under independent validations were also low, which could be a consequence of the genetic relatedness between the diversity training panel and the F5 and DH winter wheat breeding test lines. Average Rogers’ genetic coefficient between the training and test populations was 0.31, indicating genetic differences among them (S6 Table). Using GEBV alone for selection was not that successful relative to the PS and the PS+GS approaches in terms of values for response. Negative *R* values were observed for almost 50% of the datasets for both GS1 and GS2. Relying exclusively on GEBV for performing selections should therefore be taken with care, as some lines predicted to have high GEBV could have low yield. Correlations between GEBV and observed yield between a year and the next growing season under cross-validations using the AMP were in general low, indicating that high GEBV sometimes do not translate to high observed phenotypic values. This is especially true when evaluating across years due to the possible effects of genotype-by-environment interactions. In the context of selecting new parental lines based on GEBV alone, it was recently observed that selecting for high FHB resistance in winter wheat was not that reliable, as only 19% of the lines (9 out of 47) correctly predicted by GEBV belong to the best 10% for FHB resistance [35]. In another study, negative GEBV for yield were observed for synthetic hexaploid spring bread wheat lines evaluated across heat and irrigated environments [36]. Selection for drought tolerance in maize using GEBV, in contrast, has resulted in rapid genetic gains and positive selection responses through using molecular markers associated with yield under drought stress [37]. While selecting lines based on GEBV alone should be considered with caution, the implementation of genomic selection in breeding programs should help increase the rate of genetic gains through a faster breeding cycle, higher selection intensity, and efficiency of genomic prediction approaches in integrating novel genetic material in wide-crosses and pre-breeding programs [38]. Moreover, GEBV can replace phenotypes if they are more predictive of true breeding values [39]. For GEBV therefore to be more relevant in the breeding program, strategies that could help increase the selection accuracy, such as using genetically related populations, utilizing optimal training population composition and sizes, and employing ideal number of markers for predictions [40–42] should be implemented. Altogether, our results demonstrated that GEBV could still nonetheless be used as a selection criterion for grain yield in winter wheat breeding.

Selection responses achieved by integrating GWAS-derived markers as fixed effects in the prediction model (GS2) was not significantly different than that of a standard GS approach (GS1), although 17% improvement in the mean *R* was observed. This demonstrated the potential to increase gains by incorporating fixed effect markers in the model, consistent with previous studies [43,44]. It should be noted that the markers used as fixed effects in the selection model were identified to be significant only in the training population (AMP) to disregard the effect of “inside trading,” which was previously observed to cause overestimated accuracies for FHB resistance in wheat [33]. These inflated accuracies under “inside trading” are attributed to the bias caused by using significant markers that were identified in the same group of lines used for genomic predictions [33]. Using simulations, Bernardo [45] previously showed that incorporating markers with *R*^*2*^ greater than 10% in the model should give an advantage in increasing the accuracy. In the present study, significant loci with *R*^*2*^ greater than 10% were not identified. Nevertheless, even if it were the case, we still observed a positive effect of including significant markers on the predictive ability for grain yield. In addition to using GWAS-derived markers for prediction, the inclusion of genetically correlated, highly heritable traits from high-throughput field phenotyping in the prediction model have been observed to improve selection accuracy for grain yield in wheat [46–49].

Negative responses were observed for marker selection (MS) for wheat breeding lines using independent SNPs identified from association mapping using the AMP, indicating the inefficiency of using this approach exclusively for the selection of grain yield. Further, there were no LND17_F5 lines having favorable allele combinations for the most significant yield-related SNP loci, which demonstrates the difficulty of performing selections based on an MS approach (Table 2; S7 Table). In the context of genomic predictions for FHB related traits in wheat, the use of independent SNPs (i.e. markers identified using a different mapping population) was previously observed to have neutral or reducing effects on selection accuracy [33]. Marker-assisted validation, marker-aided backcrossing, and marker-assisted gene pyramiding, nonetheless, has been successfully implemented for different traits such as leaf rust resistance, powdery mildew resistance, and pre-harvest sprouting tolerance, to name a few [50]. Improvement for grain yield using MS approaches remains a challenge due to its genetic complexity, heritability, and the effects of genotype-by-environment interactions compared to disease resistance traits which are controlled by relatively few QTL with major effects [51]. Consequently, there is a need to validate results from association studies for complex traits such as yield to better implement MS strategies in the breeding program. Previously, some QTL validation studies for grain yield in wheat showed the potential of using allele specific assays such as KASP^®^ [52] to select for lines with high yield potential. Lozada et al. [53], for instance, developed marker assays for yield and component traits and used a diverse panel of spring wheat lines from CIMMYT, Mexico to validate the effects of yield-related loci previously identified in southern US winter wheat. They eventually showed the potential of developing molecular marker assays that could select for spring wheat lines with improved yield potential. In the present study, using MS alone might not necessarily result to improved gains, however, when implemented together with GS and PS approaches, improved gains in the breeding program could be observed.

## Conclusions

Gains in terms of response to selection *R*, which could be achieved by employing different selection strategies for grain yield in a winter wheat breeding program were compared. Phenotypic selection (PS) showed favorable responses to selection compared to genomic (GS) and marker selection (MS) approaches. Combining PS with GS showed a great potential in achieving higher *R* values compared to using either method alone. We observed that GS when combined with traditional PS for yield, should facilitate an increased response to selection and ultimately genetic gains in the WSU winter wheat breeding program. Altogether, we showed that genetic gains in terms of response to selection could be achieved through the integration of one selection method with another. Breeders could make important selection decisions based on the combination of one or more strategies to achieve optimal gains in plant breeding programs. Careful consideration on which selection strategies to implement, depending on the traits being evaluated, cost, and available resources should facilitate improved genetic gains for complex traits.

## Acknowledgments

The authors would like to thank Gary Shelton and Kyall Hagemeyer for assistance in the collection of yield data.

## Supporting information

**S1 Table. Genomic estimated breeding values (GEBV) for the F5 and DH winter wheat breeding lines under a standard genomic selection (GS1) scenario.**

**S2 Table. Genomic estimated breeding values (GEBV) for the F5 and DH winter wheat breeding lines with GWAS-derived markers included as fixed effects in the prediction model (GS2).**

**S3 Table. Response to selection, *R* based on phenotypic selection (PS) and marker-based selection (MS) for grain yield in US Pacific Northwest winter wheat. *R* calculated relative to the mean of the check lines.**

**S4 Table. Response to selection, *R* for GEBV-based selection (GS1 and GS2) strategies for grain yield in US Pacific Northwest winter wheat. *R* calculated relative to the mean of the check lines.**

**S5 Table. Response to selection, *R* for phenotypic + genomic (PS+GS1 and PS+GS2) selection strategies and the number of lines selected in combining both approaches for selection (in parentheses) of yield in US Pacific Northwest winter wheat. *R* calculated relative to the mean of the check lines.**

**S6 Table. Roger’s genetic coefficient between the association mapping training panel (AMP) and the winter wheat breeding lines across each chromosome.**

**S7 Table. Winter wheat breeding lines selected under a marker-based selection (MS) strategy using the five most significant SNPs identified using a GWAS approach for a diverse winter wheat mapping panel.**

**S1 File. Genotype data (16,233 SNP markers) for the winter wheat association mapping panel (AMP) used for genomewide association study.**

**S2 File. Genotype data (11,089 SNP markers) for the winter wheat breeding lines used for genomic predictions for grain yield. This panel is a subset of the 16,233 markers used for the AMP (S1 File).**

**S3 File. Adjusted yield (t/ha) for each site-year combination for the winter wheat association mapping panel (AMP).**

**S4 File. Adjusted yield (t/ha) for the F5 and DH winter wheat breeding lines.**

## References

1. Hickey LT, N. Hafeez A, Robinson H, Jackson SA, Leal-Bertioli SCM, Tester M, et al. Breeding crops to feed 10 billion. Nat Biotechnol. 2019; Doi:10.1038/s41587-019-0152-9

2. Moose SP, Mumm RH. Molecular plant breeding as the foundation for 21st century crop improvement. Plant Physiol. 2008;147: 969–977.

3. Cobb JN, Juma RU, Biswas PS, Arbelaez JD, Rutkoski J, Atlin G, et al. Enhancing the rate of genetic gain in public-sector plant breeding programs: lessons from the breeder’s equation. Theor Appl Genet. 2019;132: 627–645. Doi:10.1007/s00122-019-03317-0

4. Li H, Rasheed A, Hickey LT, He Z. Fast-Forwarding Genetic Gain. Trends Plant Sci. 2018;23: 184–186. Doi:https://doi.org/10.1016/j.tplants.2018.01.007

5. Simons KJ, Fellers JP, Trick HN, Zhang Z, Tai Y-S, Gill BS, et al. Molecular Characterization of the Major Wheat Domestication Gene <em>Q</em>; Genetics. 2006;172: 547 LP–555. Doi:10.1534/genetics.105.044727

6. Venske E, dos Santos RS, Busanello C, Gustafson P, de Oliveira A. Bread wheat: a role model for plant domestication and breeding. Hereditas. 2019;156: 16. Doi:10.1186/s41065-019-0093-9

7. Rutkoski J, Singh RP, Huerta-Espino J, Bhavani S, Poland J, Jannink JL, et al. Genetic Gain from Phenotypic and Genomic Selection for Quantitative Resistance to Stem Rust of Wheat. Plant Genome. 2015;8. Doi:10.3835/plantgenome2014.10.0074

8. Anderson JA, Chao S, Liu S. Molecular Breeding Using a Major QTL for Fusarium Head Blight Resistance in Wheat. Crop Sci. 2007;47: S-112-S-119. Doi:10.2135/cropsci2007.04.0006IPBS

9. Miedaner T, Wilde F, Korzun V, Ebmeyer E, Schmolke M, Hartl L, et al. Marker selection for Fusarium head blight resistance based on quantitative trait loci (QTL) from two European sources compared to phenotypic selection in winter wheat. Euphytica. 2009;166: 219–227. Doi:10.1007/s10681-008-9832-0

10. Wilde F, Korzun V, Ebmeyer E, Geiger HH, Miedaner T. Comparison of phenotypic and marker-based selection for Fusarium head blight resistance and DON content in spring wheat. Mol Breed. 2007;19: 357–370. Doi:10.1007/s11032-006-9067-5

11. Quarrie SA, Steed A, Calestani C, Semikhodskii A, Lebreton C, Chinoy C, et al. A high-density genetic map of hexaploid wheat (Triticum aestivum L.) from the cross Chinese Spring {\texttimes} SQ1 and its use to compare QTLs for grain yield across a range of environments. Theor Appl Genet. 2005;110: 865–880. Doi:10.1007/s00122-004-1902-7

12. Li F, Wen W, Liu J, Zhang Y, Cao S, He Z, et al. Genetic architecture of grain yield in bread wheat based on genome-wide association studies. BMC Plant Biol. 2019;19: 168. Doi:10.1186/s12870-019-1781-3

13. Garcia M, Eckermann P, Haefele S, Satija S, Sznajder B, Timmins A, et al. Genome-wide association mapping of grain yield in a diverse collection of spring wheat (Triticum aestivum L.) evaluated in southern Australia. PLoS One. 2019;14: e0211730. Available: https://doi.org/10.1371/journal.pone.0211730

14. Green AJ, Berger G, Griffey CA, Pitman R, Thomason W, Balota M, et al. Genetic yield improvement in soft red winter wheat in the Eastern United States from 1919 to 2009. Crop Sci. 2012;52: 2097–2108.

15. Rodríguez F, Alvarado G, Pacheco Á, Burgueño J. ACBD-R. Augmented Complete Block Design with R for Windows. Version 4.0 [Internet]. CIMMYT Research Data & Software Repository Network; 2018. Doi:11529/10855

16. Peterson CJ, Allan RE, Rubenthaler GL, Line RF. Registration of ‘Eltan’ Wheat. Crop Sci. 1991;31: 1704. Doi:10.2135/cropsci1991.0011183X003100060075x

17. Allan RE, Peterson CJ, Rubenthaler GL, Line RF, Roberts DE. Registration of ‘Madsen’wheat. Crop Sci. 1989;29: 1575–1576.

18. Jones SS, Murray TD, Lyon SR, Morris CF, Line RF. Registration ofBruehl’wheat.(Registrations of Cultivars). Crop Sci. 2001;41: 2006–2008.

19. Carter AH, Jones SS, Lyon SR, Balow KA, Shelton GB, Higginbotham RW, et al. Registration of ‘Otto’wheat. J Plant Regist. 2013;7: 195–200.

20. Carter AH, Jones SS, Balow KA, Shelton GB, Burke AB, Lyon S, et al. Registration of ‘Jasper’soft white winter wheat. J Plant Regist. 2017;11: 263–268.

21. Jones SS, Lyon SR, Balow KA, Gollnick MA, Murphy KM, Kuehner JS, et al. Registration of ‘Xerpha’wheat. J plant Regist. 2010;4: 137–140.

22. Zemetra RS, Souza EJ, Lauver M, Windes J, Guy SO, Brown B, et al. Registration of ‘Brundage’wheat. Crop Sci. 1998;38: 67.

23. Carter AH, Jones SS, Cai X, Lyon SR, Balow KA, Shelton GB, et al. Registration of ‘Puma’soft white winter wheat. J Plant Regist. 2014;8: 273–278.

24. Lozada DN, Carter AH. Accuracy of single and multi-trait genomic prediction models for grain yield in US Pacific Northwest winter wheat. 2019. Submitted. Crop Breeding, Genetics, and Genomics.

25. Poland, Jesse; Endelman, Jeffrey; Dawson, Julie; Rutkoski, Jessica; Wu, Shuangye; Manes, Yann; Dreisigacker, Susanne; Crossa, Jose and Sanchez-Villeda, Hector and Sorrells M and others. Genomic selection in wheat breeding using genotyping-by-sequencing. Plant Genome. 2012;5.

26. Liu X, Huang M, Fan B, Buckler ES, Zhang Z. Iterative Usage of Fixed and Random Effect Models for Powerful and Efficient Genome-Wide Association Studies. PLOS Genet. 2016;12: e1005767. Available: https://doi.org/10.1371/journal.pgen.1005767

27. R Development Core Team. R: A Language and Environment for Statistical Computing. 2018. Vienna, Austria.

28. Benjamini Y, Hochberg Y. Controlling the false discovery rate: a practical and powerful approach to multiple testing. J R Stat Soc Ser B. 1995; 289–300.

29. SAS Institute. SAS System Options: Reference, 2nd ed. Cary, NC: SAS Institute; 2015.

30. Chen CJ, Zhang Z. iPat: intelligent prediction and association tool for genomic research. Bioinformatics. 2018;34: 1925–1927. Available: http://dx.doi.org/10.1093/bioinformatics/bty015

31. Endelman JB. Ridge regression and other kernels for genomic selection with R package rrBLUP. Plant Genome. 2011;4: 250–255.

32. Falconer DS. Introduction to Quantitative Genetics. 3rd ed. New York: Longman Scientific and Technical; 1989.

33. Arruda MP, Lipka AE, Brown PJ, Krill AM, Thurber C, Brown-Guedira G, et al. Comparing genomic selection and marker-assisted selection for Fusarium head blight resistance in wheat (Triticum aestivum L.). Mol Breed. 2016;36: 84. Doi:10.1007/s11032-016-0508-5

34. Belamkar V, Guttieri MJ, Hussain W, Jarquín D, El-basyoni I, Poland J, et al. Genomic Selection in Preliminary Yield Trials in a Winter Wheat Breeding Program. G3 Genes|Genomes|Genetics. 2018;8: 2735 LP–2747. Available: http://www.g3journal.org/content/8/8/2735.abstract

35. Herter CP, Ebmeyer E, Kollers S, Korzun V, Miedaner T. An experimental approach for estimating the genomic selection advantage for Fusarium head blight and Septoria tritici blotch in winter wheat. Theor Appl Genet. 2019; Doi:10.1007/s00122-019-03364-7

36. Jafarzadeh J, Bonnett D, Jannink J-L, Akdemir D, Dreisigacker S, Sorrells ME. Breeding Value of Primary Synthetic Wheat Genotypes for Grain Yield. PLoS One. 2016;11: e0162860. Available: https://doi.org/10.1371/journal.pone.0162860

37. Vivek BS, Krishna GK, Vengadessan V, Babu R, Zaidi PH, Kha LQ, et al. Use of Genomic Estimated Breeding Values Results in Rapid Genetic Gains for Drought Tolerance in Maize. Plant Genome. 2017;10. Doi:10.3835/plantgenome2016.07.0070

38. Hickey JM, Chiurugwi T, Mackay I, Powell W, Participants IGS in CBPW, Hickey JM, et al. Genomic prediction unifies animal and plant breeding programs to form platforms for biological discovery. Nat Genet. 2017;49: 1297. Available: https://doi.org/10.1038/ng.3920

39. Lorenz AJ. Resource Allocation for Maximizing Prediction Accuracy and Genetic Gain of Genomic Selection in Plant Breeding: A Simulation Experiment. G3 Genes|Genomes|Genetics. 2013;3: 481 LP–491. Doi:10.1534/g3.112.004911

40. Zhong S, Dekkers JCM, Fernando RL, Jannink J-L. Factors Affecting Accuracy From Genomic Selection in Populations Derived From Multiple Inbred Lines: A Barley Case Study. Genetics. 2009;182: 355–364. Doi:10.1534/genetics.108.098277

41. Spindel J, Begum H, Akdemir D, Virk P, Collard B, Redoña E, et al. Genomic Selection and Association Mapping in Rice (Oryza sativa): Effect of Trait Genetic Architecture, Training Population Composition, Marker Number and Statistical Model on Accuracy of Rice Genomic Selection in Elite, Tropical Rice Breeding Lines. PLOS Genet. 2015;11: 1–25. Doi:10.1371/journal.pgen.1004982

42. Cericola F, Jahoor A, Orabi J, Andersen JR, Janss LL, Jensen J. Optimizing Training Population Size and Genotyping Strategy for Genomic Prediction Using Association Study Results and Pedigree Information. A Case of Study in Advanced Wheat Breeding Lines. PLoS One. 2017;12: e0169606. Available: https://doi.org/10.1371/journal.pone.0169606

43. Michel S, Kummer C, Gallee M, Hellinger J, Ametz C, Akgöl B, et al. Improving the baking quality of bread wheat by genomic selection in early generations. Theor Appl Genet. 2018;131: 477–493. Doi:10.1007/s00122-017-2998-x

44. Galiano-Carneiro AL, Boeven PHG, Maurer HP, Würschum T, Miedaner T. Genome-wide association study for an efficient selection of Fusarium head blight resistance in winter triticale. Euphytica. 2018;215: 4. Doi:10.1007/s10681-018-2327-8

45. Bernardo R. Genomewide Selection when Major Genes Are Known. Crop Sci. 2014;54: 68–75. Doi:10.2135/cropsci2013.05.0315

46. Rutkoski J, Poland J, Mondal S, Autrique E, Pérez LG, Crossa J, et al. Canopy Temperature and Vegetation Indices from High-Throughput Phenotyping Improve Accuracy of Pedigree and Genomic Selection for Grain Yield in Wheat. G3 (Bethesda). 2016;6: 2799–2808. Doi:10.1534/g3.116.032888

47. Crain J, Mondal S, Rutkoski J, Singh RP, Poland J. Combining High-Throughput Phenotyping and Genomic Information to Increase Prediction and Selection Accuracy in Wheat Breeding. Plant Genome. 2018;11. Doi:10.3835/plantgenome2017.05.0043

48. Sun J, Poland JA, Mondal S, Crossa J, Juliana P, Singh RP, et al. High-throughput phenotyping platforms enhance genomic selection for wheat grain yield across populations and cycles in early stage. Theor Appl Genet. 2019; Doi:10.1007/s00122-019-03309-0

49. Lozada DN, Godoy J V, Carter AH. Genomic prediction and indirect selection for grain yield using spectral reflectance indices from high-throughput phenotyping. 2019. In prep.

50. 1 102 Gupta PK, Langridge P, Mir RR. Marker-assisted wheat breeding: present status and future possibilities. Mol Breed. 2010;26: 145–161. Doi:10.1007/s11032-009-9359-7

51. Hospital F. Challenges for effective marker-assisted selection in plants. Genetica. 2009;136: 303–310. Doi:10.1007/s10709-008-9307-1

52. Semagn K, Babu R, Hearne S, Olsen M. Single nucleotide polymorphism genotyping using Kompetitive Allele Specific PCR (KASP): overview of the technology and its application in crop improvement. Mol Breed. 2014;33: 1–14. Doi:10.1007/s11032-013-9917-x

53. Lozada DN, Mason RE, Sukumaran S, Dreisigacker S. Validation of grain yield QTLs from soft winter wheat using a CIMMYT spring wheat panel. Crop Sci. 2018;58:1964–1971. Doi:10.2135/cropsci2018.04.0232

